# Dimerization of the Pragmin pseudo-kinase regulates protein tyrosine phosphorylation

**DOI:** 10.1101/218669

**Authors:** Céline Lecointre, Valérie simon, Clément kerneur, Frédéric Allemand, Aurélie Fournet, Ingrid Montarras, Jean-Luc Pons, Muriel Gelin, Constance Brignatz, Serge Urbach, Gilles Labesse, Serge Roche

## Abstract

The pseudo-kinase and signaling protein Pragmin has been linked to cancer by regulating protein tyrosine phosphorylation via unknown mechanisms. Here we present the crystal structure of the Pragmin 906-1368 amino acids C-terminus, which encompasses its kinase domain. We show that Pragmin contains a classical protein kinase fold devoid of catalytic activity. A particular inhibitory triad, conserved in a Pragmin/SgK269/PEAK1/C19orf35 superfamily, tightly holds the catalytic lysine (K997) to prevent ATP binding. By proteomics, we discovered that this pseudo-kinase uses the tyrosine kinase CSK to induce protein tyrosine phosphorylation in human cells. Interestingly, the protein kinase domain is bordered by N- and C-terminal extensions forming an original dimerization domain that regulates Pragmin self-association and stimulates CSK activity. A1329E mutation in the C-terminal extension destabilizes Pragmin dimerization and reduces CSK activation. Thus, our results reveal a new dimerization mechanism by which a pseudo-kinase can induce protein tyrosine phosphorylation.

## INTRODUCTION

Protein phosphorylation plays a pivotal role in cell biological events that leads to cell growth, adhesion, survival or differentiation (Hunter, 2000). Deregulation of protein kinases plays important roles in human diseases such as cancer (Blume-Jensen & Hunter, 2001). As results, they have become highly attractive therapeutic targets and several small inhibitors are currently used in the clinic. However, this strategy might not apply to the whole human kinome, as it contains about 50 pseudo-kinases out of roughly 520 protein kinases, which are predicted to be catalytically inactive due to the lack of important residues required for full enzymatic activity (Boudeau et al, 2006; Murphy et al, 2014). Little information is available on the precise role of these pseudo-kinases in cellular signaling however recent structural analyses indicate that they function by docking to and stimulating catalytically active protein kinases for efficient protein phosphorylation and that some of them, after all, possess a weak active kinase activity (Mukherjee et al, 2008), thus implicating an unconventional molecular mechanism of protein phosphorylation (Jacobsen & Murphy, 2017).

Recent reports have linked pseudo-kinases, such as human Pragmin, and cancer. Pragmin, also known as SgK223, was originally identified as an effector of the small GTPase Rnd2 from a rat expression library and a novel regulator of cell adhesion and morphology(Tanaka et al, 2006). Pragmin belongs to a subfamily of pseudo-kinases that include SgK269/PEAK1 involved in adhesive and growth factor receptor signaling leading to cell growth and adhesion (Croucher et al, 2013; Kelber et al, 2012; Rozakis-Adcock et al, 1992; Wang et al, 2010). In neuronal cells, Pragmin induces Rho-dependent cell contraction and negatively regulates neurite outgrowth (Tanaka et al, 2006). In cancer, human Pragmin is overexpressed in various adenocarcinoma cells and promotes cell migration and invasion (Leroy et al, 2009; Senda et al, 2016; Tactacan et al, 2015). This invasive function of Pragmin is independent from MAPKs and AKT activities but implicates a Jak1/STAT3-dependent mechanism in pancreatic tumor cells (Tactacan et al, 2015). In gastric adenocarcinoma cells, Pragmin expression induces cell migration by a CSK-dependent mechanism (Senda et al, 2016). By comprehensive phosphoproteomic analyses of metastatic colorectal cancer (CRC) cells, we have identified human Pragmin as a critical component of oncogenic signaling driven by the tyrosine kinase (TK) Src to induce tumor cell growth and invasion. Pragmin also mediates protein tyrosine phosphorylation in these cancer cells, indicating that this signaling protein could exert its tumoral function by a novel TK-dependent mechanism to be characterized (Leroy et al, 2009; Sirvent et al, 2015; Sirvent et al, 2012b). Accordingly, Pragmin possesses a protein kinase domain at the C-terminus, suggesting that it might have an active protein kinase activity as reported for SgK269/PEAK1 (Wang et al, 2010). However, conserved sequences required for catalytic reaction, such as the Gly-rich loop and the so-called DFG motif important for Mg2+ binding are missing. Thus, the exact role of the Pragmin kinase domain in cell signaling and malignant cell transformation needs to be clarified. Here we have addressed this question by a structural analysis of Pragmin C-terminus encompassing its protein kinase domain. We show that Pragmin contains a classical protein kinase fold devoid of catalytic activity due to the presence of a particular inhibitory triad that prevents ATP binding to the catalytic lysine (K997). We also reveal that the pseudo-kinase domain is bordered by two original N- and C-terminal extensions that regulate Pragmin self-association in human cells. Biochemical analysis of Pragmin expression in human cells confirms our structural model and proteomics identified CSK as the TK mediating Pragmin-induced protein tyrosine phosphorylation. Finally, by taking advantage of our structural data, we propose a model in which protein homo-dimerization regulates Pragmin protein tyrosine phosphorylation.

## RESULTS

### Overall Structure and topology of Pragmin kinase domain

We first delineated the Pragmin kinase domain for crystallographic analysis (Supplementary Fig S1). The C-terminal region of Pragmin shares strong similarities to that of Sgk269/PEAK1 but theses sequences significantly differ from sequences of conventional protein kinases. In order to refine the domain boundaries and to define a stable and soluble domain to be over-expressed, a thorough analysis of multiple sequence-structure alignments was performed using the program ViTO (Catherinot & Labesse, 2004). This analysis led to a putative delimitation of a complete protein kinase domain (residues 950 to 1292), despite the divergence and the absence of various important motifs for enzymatic activity (Supplementary Fig S1). In addition, sequence conservation among the Pragmin/Sgk269/PEAK1 family prompted us to include the whole very C-terminal segment (1293-1368) as well as a short N-terminal extension. Among four constructs tested, the longest one (Pragmin 906-1368, hereafter, named Pragmin C) appeared well expressed in *E coli* and stable after purification. The structure of this purified domain was next solved by molecular replacement despite the low overall sequence similarity (20-25%) detected with known protein kinases, by taking advantage of a structure-ensemble computed from 10 distinct templates, as a search model to increased signal-to-noise-ratio (Trapani et al, 2006). The crystal structure was refined to a Rwork/Rfree of 19.0/24.3 at a 2.8 Å resolution with two molecules in the asymmetric unit (Supplementary Table S1). The two molecules are highly similar (RMSD ~0.86 Å over 330 residues) despite changes in their crystal packing environment (Fig 1). They mainly differ by some loops partially visible in one and not the other monomer or some local conformational rearrangements. In total, almost 82 residues cannot be modeled in either monomer (102 in each monomer) of the refined structure. These regions mostly correspond to variable segments showing highly variable sequences and amino acid composition bias (Supplementary Fig S1). Accordingly they are predicted to be mainly unfolded or natively disordered in agreement with the slightly more negative signal at 208 nm compared to 222 nm in the UV-CD spectrum (Supplementary Fig S2). The folded part of the Pragmin C is a monomer of approximate dimensions 70 × 60 × 40 Å. Each of the two independent monomers face a crystal mate through an interface built up by four long helices from the N-terminal and C-terminal extension (see below). This interface is huge (~4180 squared Å), suggesting that Pragmin dimers are very stable in solution even at very low concentration. Indeed, Pragmin C was stable and perfectly dimeric in solution, according to MALLS, DLS, and SAXS studies at low micromolar concentrations (data not shown). The overall fold of Pragmin kinase domain is clearly but distantly related to the conserved fold of conventional protein kinases. A common core was delineated by FATCAT showing a RMSD of ~3 Å over ~220 residues with a mean sequence identity ~13%. The N-terminal kinase domain (residues 950-1071) is mainly composed of 5 β-strands. The kinase C-terminal region (residues 1072-1292) is mainly composed of α-helices (labelled D to H according to the PK nomenclature) showing the same arrangements seen in other protein kinases. The protein kinase C-lobe possesses variant sequences corresponding to Hanks motifs VII-XI, while motifs I-VIa of the N-lobe including the Glycine-rich loop were highly degenerated; one exception was the catalytic lysine that appeared to be conserved (Supplementary Fig S1). Overall, these results indicate that the core of the Pragmin protein kinase fold displays common features related to conventional protein kinases.

**Figure 1:**
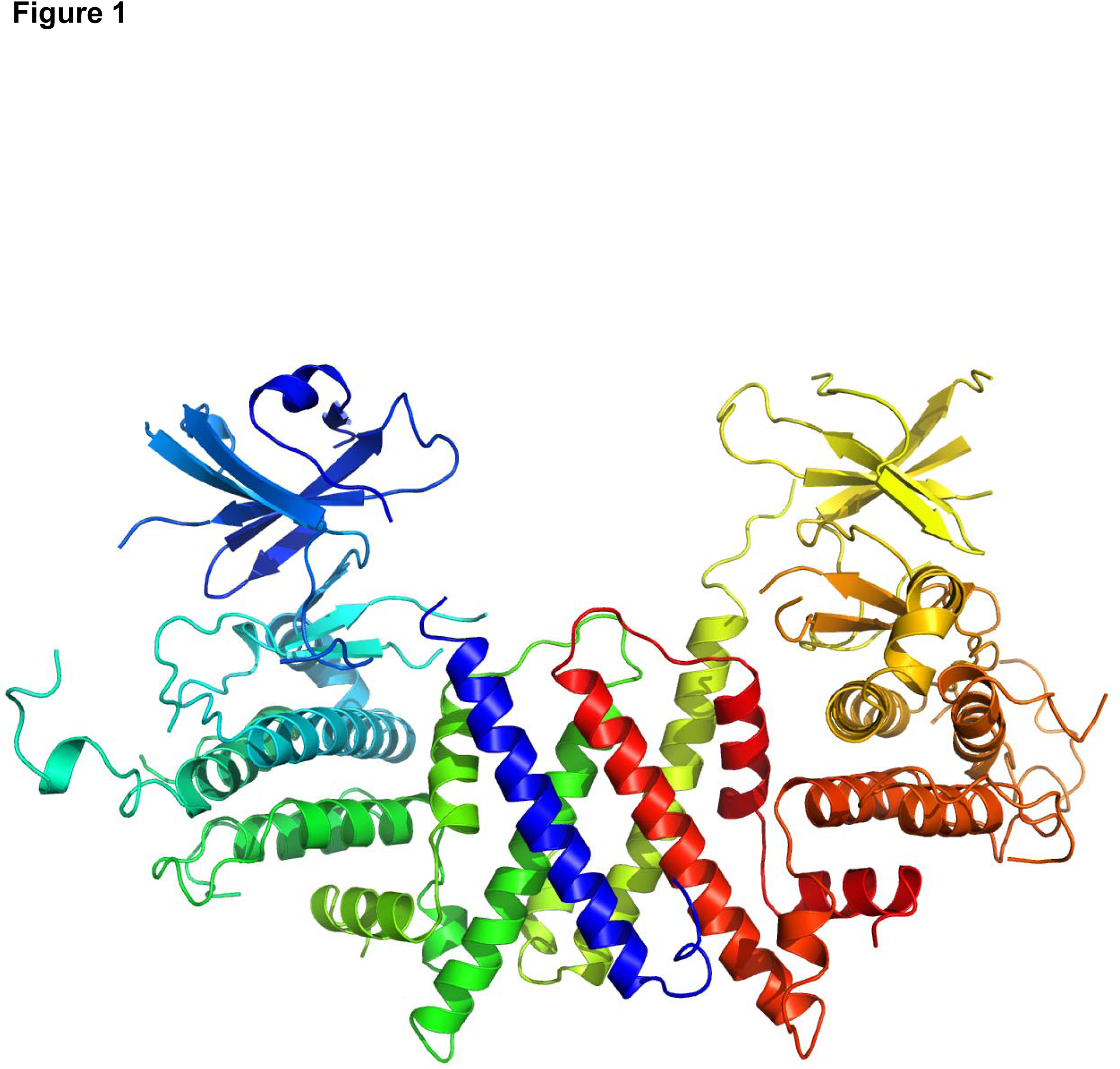
**Overall structure and topology of Pragmin C-terminus.** It includes a protein kinase domain and a dimerization module. Ribbon diagram of the two monomers. The color rainbow runs from blue at the N-terminus of a monomer to red at the C-terminus of the second monomer. Picture drawn using Pymol.

### Pragmin is a pseudo-kinase

Further scrutiny of the crystal structure suggests that Pragmin does not possess any ATP-binding activity. First of all, it seems to lack any cavity at the interface between the Nand C-lobes corresponding the ATP-binding cleft, which is conserved in all enzymatically active protein kinases. Indeed, each independent monomer of the current crystal structure adopts a tightly closed conformation, which would prevent the entrance of any nucleotide, despite their distinctive crystal packing environment and the presence of ADP (10 mM) in the crystallization media (Fig 2A and B). In addition, residues lying this cavity show significant and particular substitutions suggesting disruption of the ATP-binding properties. For instance, the DFG motif is replaced by a NFL sequence and the glycine-rich loop contains only one glycine. More importantly, three residues, which were highly conserved among most members of the family (see below), seem to hijack the lysine. Indeed, the aspartate D978, the tyrosine Y981 and the glutamine Q1021 form an intricate hydrogen bond network with the sidechain amino group of K997 (Fig 2A and B). This tight interactions make the lysine completely inaccessible to any nucleotide and may form an 'inhibitory triad' to prevent catalysis. Consistent with a lack of predicted enzymatic activity, the canonical Hanks motifs appear poorly conserved among Pragmin and SgK269/PEAK1 sequences (Supplementary Fig S1). No ATP binding was detected for the recombinant domain or its variant (D978N, Y981F or Q1021E) when assessed by thermal shift assay (Fig 2B), while all mutants were properly folded as the wild-type protein (Supplementary Fig S2). Furthermore, adenosine derivatives harboring cysteine-reactive function failed to react with the neighboring cysteine C999, unlike the active TK Src (data not shown). Finally, Pragmin expression in human HEK293T cells did not show specific ATP binding activity in contrast to Src, when assessed for its avidity to the destobiotin-ATP analog (Fig 2c). Collectively, this data indicate that Pragmin is indeed a pseudo-kinase.

**Figure 2:**
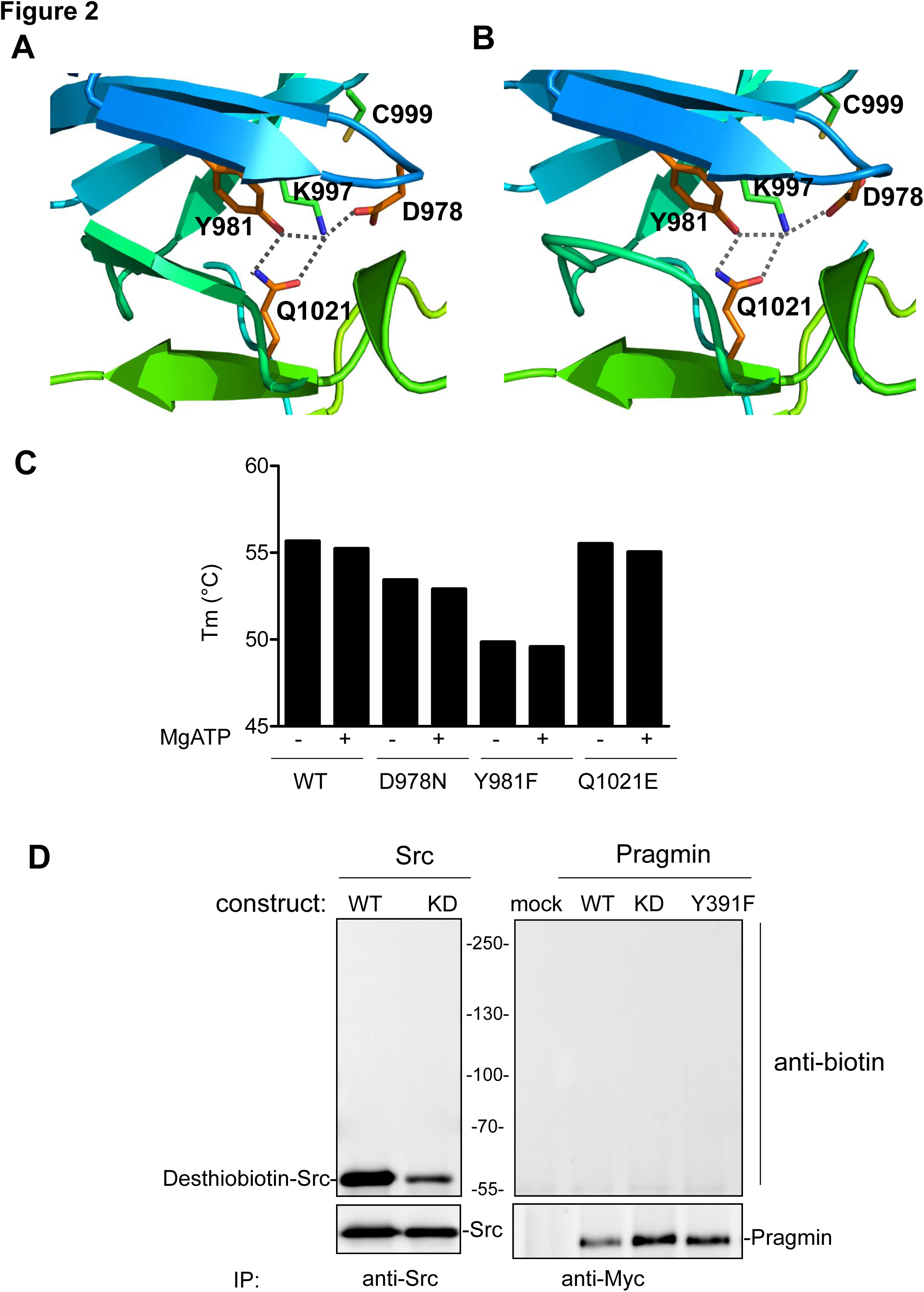
**Pragmin is a pseudo-kinase. A.** Close view of the putative ATP binding site in monomer A. Main chain is shown as in Figure 1, while side-chains of the conserved lysine K997 and the neighboring residues (see text) are shown in wireframe. The aspartate D978, the tyrosine Y981 and the glutamine Q1021 form an intricate network of hydrogen bonds (in dashed and grey lines) with the lysine K997, which is predicted to prevent ATP binding. **B.** Same as A in monomer B. **C.** Absence of thermostabilisation of wild-type and mutated Pragmin by Mg-ATP. Wild-type and the three mutated proteins (at 2 µM final concentration) were submitted to thermal shift assays in absence or presence of 10 mM ATP (identical results were obtained at 5 mM; not shown). No significant shift were detected for the recombinant enzyme or any mutant by addition of the nucleotide. In parallel, some destabilization appeared induced by the mutations as observed for two mutants D978N (-2.2 °C) and Y981F (-5.8 °C). **D.** Exogenous Pragmin does not bind to ATP-desthobiotin in HEK293T cells, unlike Src. Is shown the level of desthobiotinated proteins from immunoprecipitated Pragmin and Src (wild-type and mutated) proteins that were expressed in HEK293T cells as shown. The level of immunoprecipitated Pragmin and Src proteins is also shown. Picture drawn using Pymol.

### An original dimerization module for a scaffolding activity

Our structural data also reveal that the first forty-four residues (906-949) and the last seventy-six residues (1293-1368) form a five-helix module, which is involved in the dimerization of two crystal-mate monomers (Fig 3). This additional module appears highly original with no similarity with already known structures. It is composed of two long helices (921-948 and 1306-1334) and three shorter ones (911-919, 1341-1353 and 1355-1366). The latter three seem to organize the positioning relative to the pseudo-kinase domain, of the two long helices that directly participate in the dimerization interface. While four-helix bundles are quite frequent in dimerization modules, they usually arrange in a parallel or anti-parallel manner. An original organization dimerization was observed in the case of Pragmin (Fig 3B). Indeed, the long dimerization helix from the N-terminal region interacts with the long dimerization helix from the very C-terminal region with a ~40° tilt to form a crossing sign or a 'X' symbol. The dimerization is built by two tightly contacting crossings, with the N-terminal helix from one monomer being parallel to the C-terminal one from the second monomer (Fig 3). The three other helical segments of the dimerization module appear to interact with the protein kinase domain and to connect it with the dimerization helices. The huge and well-structured interface suggests that it is stable and functionally important.

**Figure 3:**
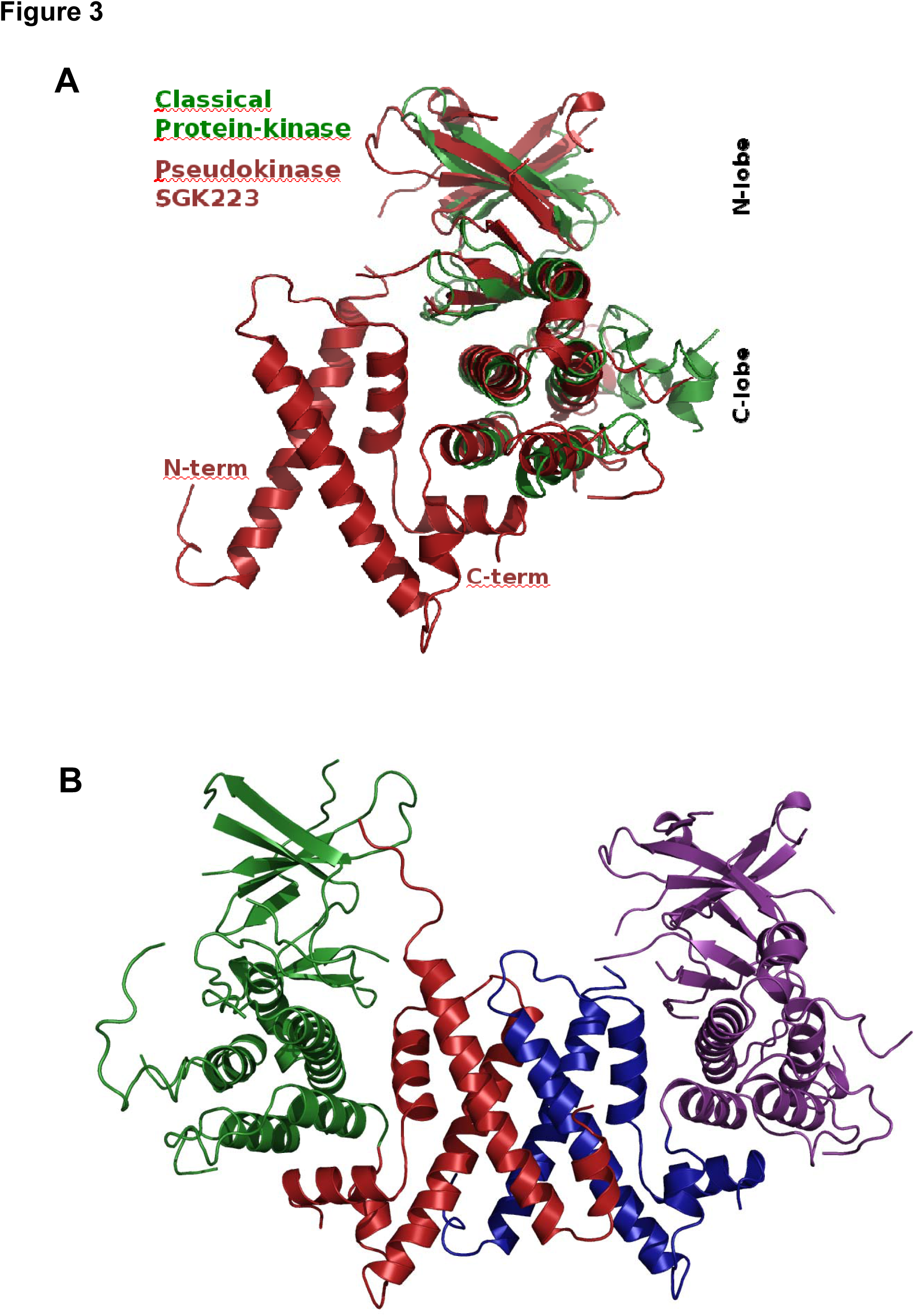
**An original dimerization module. A.** Superposition of one monomer of Pragmin (red ribbon) and a classical protein kinase in close conformation (green ribbon). **B.** Pragmin dimer built by two crystal mates. The protein kinase domains are shown in green and purple ribbons while the dimerization module are shown in red and blue ribbons.

### Conserved kinase topology in a Pragmin/Sgk269/PEAK1/C19orf35 superfamily

Sequence searches using the segment corresponding to the very C-terminal region and comprising most of the dimerization module (1293-1368) showed strong similarities with other Pragmin and SgK269/PEAK1 orthologues (Supplementary Fig S1). Importantly, they harbor conserved motifs corresponding to residues buried in this extra helical module suggesting that the same architecture is preserved in all these pseudo-kinases. Interestingly, these searches also highlighted the conservation of these motifs in a subfamily of unannotated proteins, whose human member is C19orf35. The latter also harbors a pseudo-kinase domain and shows a weak but significant level of sequence similarities with Pragmin and SgK269/PEAK1 (~29% of overall sequence identity but up to 38% in the dimerization region) including the specific inhibitory triad (see above); therefore C19orf35 may indeed be another member of the Pragmin/SgK269/PEAK1 subfamily of pseudo-kinases, sharing a common dimerization module for all of them (Fig 4). In addition, these three pseudo-kinases also contain specific and possibly intrinsically disordered N-terminal extensions, which is very short in C19orf35 (~100 residues) and much longer in Pragmin (~ 900 residues) and Sgk269/PEAK1 (~1200 residues). These specific domains may dictate specific functions for these pseudo-kinases.

**Figure 4:**
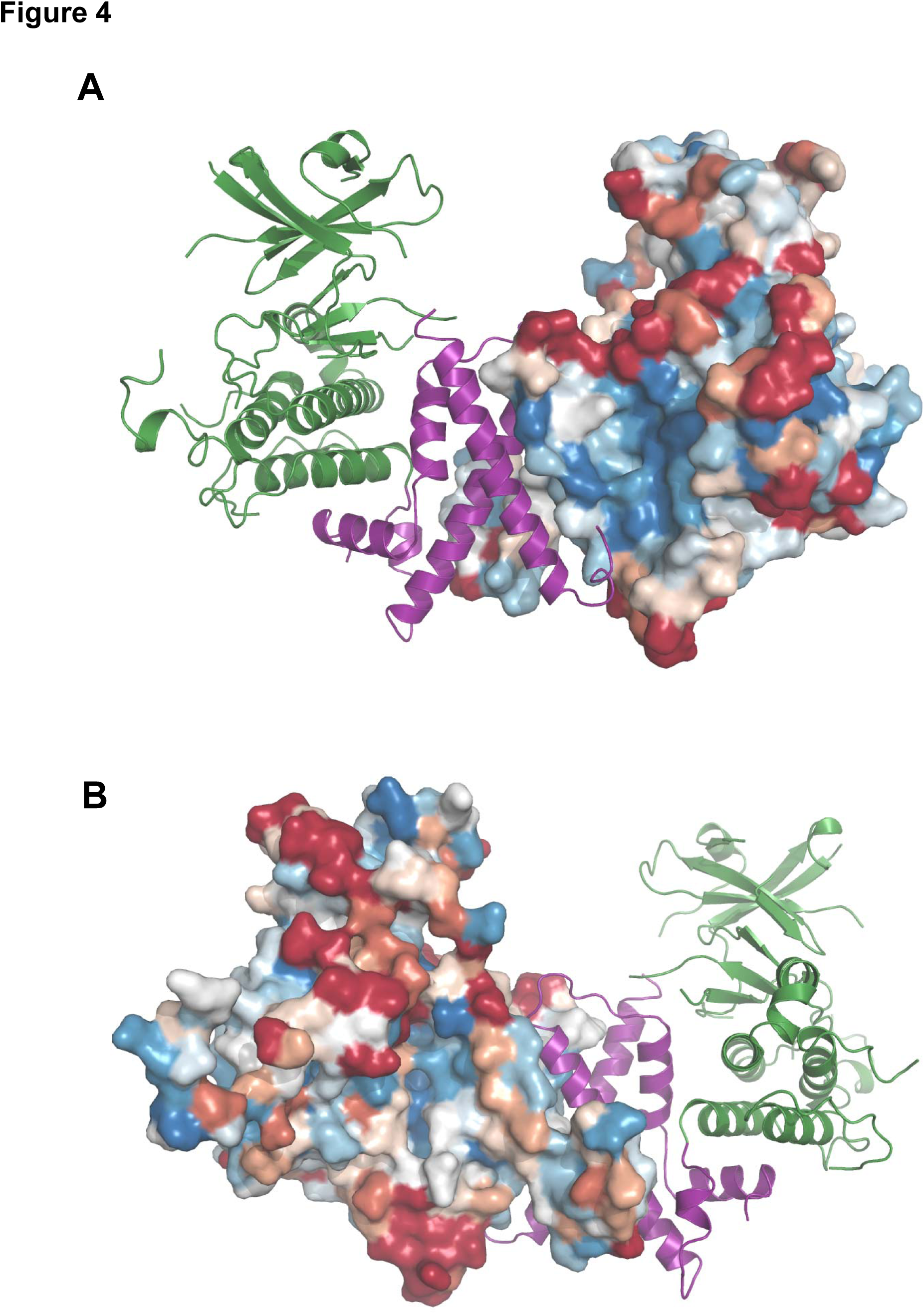
**Evolutionary trace on the Pragmin dimer. A.** Sequence conservation among Pragmin/SgK269/PEAK1/Cs035 orthologs was computing using CONSURF (Ashkenazy et al, 2016) and display as a color code (from highly conserved in blue to poorly conserved in red) on the protein surface (right monomer). The second monomer is shown as a ribbon in green (pseudo-kinase domain) and purple (dimerization module) color respectively, to highlight the position of the dimer interface. The latter appears much better conserved (light/deep blue) compared to the putative ATP-binding site. **B.** Same as in panel A but with a 180° rotation.

### Pragmin self-association in human cells

We next interrogated the biological relevance of our structural data and first investigated Pragmin self-association in human cells. A FLAG-tagged Pragmin construct was co-expressed with constructs expressing either Myc-tagged wild-type or various Pragmin mutants in HEK293T cells (Fig 5A) and Pragmin self-association was analyzed by coimmunoprecipitation. We found a large association of expressed FLAG-tagged with Myc-tagged Pragmin proteins consistent with Pragmin homo-dimerization occurring in human cells (Fig 5B). Structure-association analysis next revealed that Pragmin dimerization does not require the catalytic K997 nor the main tyrosine phosphorylation site Y391 (Leroy et al, 2009). However a Pragmin C-terminal domain mutant (857-1368) encompassing the 2 dimerization segments was sufficient to associate with wild-type Pragmin, while N-terminal Pragmin constructs lacking these domains did not. We next interrogated the contribution of each segment on Pragmin self-association by a similar method, however partial deletion of the C-terminal dimerization region (1293-1368) produced a highly unstable protein, precluding a direct assessment of this region on this molecular process (Supplementary Fig S3). We then took advantage of our crystal structure to design point mutations potentially affecting the dimerization stability (Fig 5C). Four mutations were designed to substitute a charged amino acid to hydrophobic and buried residues, with the aim of disrupting the quaternary structure while maintaining the local secondary structure. Each of the three mutation (L935E in the N-terminal and Y349E and L1369E in the C-terminal region) were insufficient to disrupt protein dimerization in HEK293T cells (Fig 5B). This may correlate to the similar size of the substituted amino acid compared to the wild-type residue. Despite the charge added in a buried and hydrophobic environment, the very large surface of that interface may compensate for the local destabilization. On the contrary, we found a strong diminution of Pragmin self-association upon A1329E mutation in the C-terminal extension (Fig 5B). This result is in agreement with the substitution of a small alanine toward glutamate, a much larger and charged residue. This dramatic change was expected to create not only some charge repulsion but also some strong van der Waals clashes (Fig 5D). Indeed, one A1329 from one monomer faces its symmetrical position A1329' from the second monomer. This key position and the larger modification were expected to be much more detrimental for the interface. Overall, these biochemical observations validate our structural model and point to a central role of the pseudo-kinase domain N and C-terminal extensions in Pragmin self-association.

**Figure 5:**
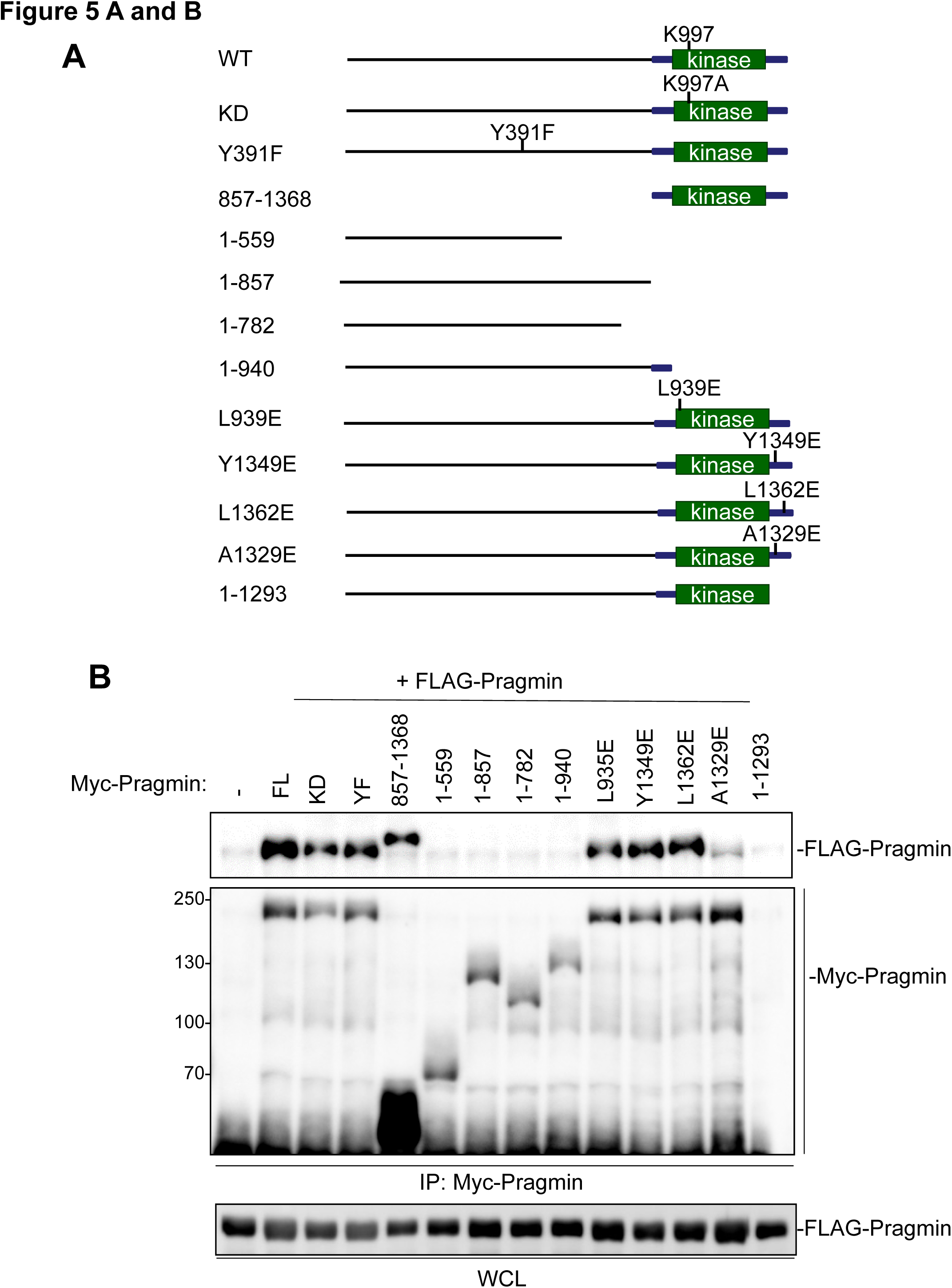
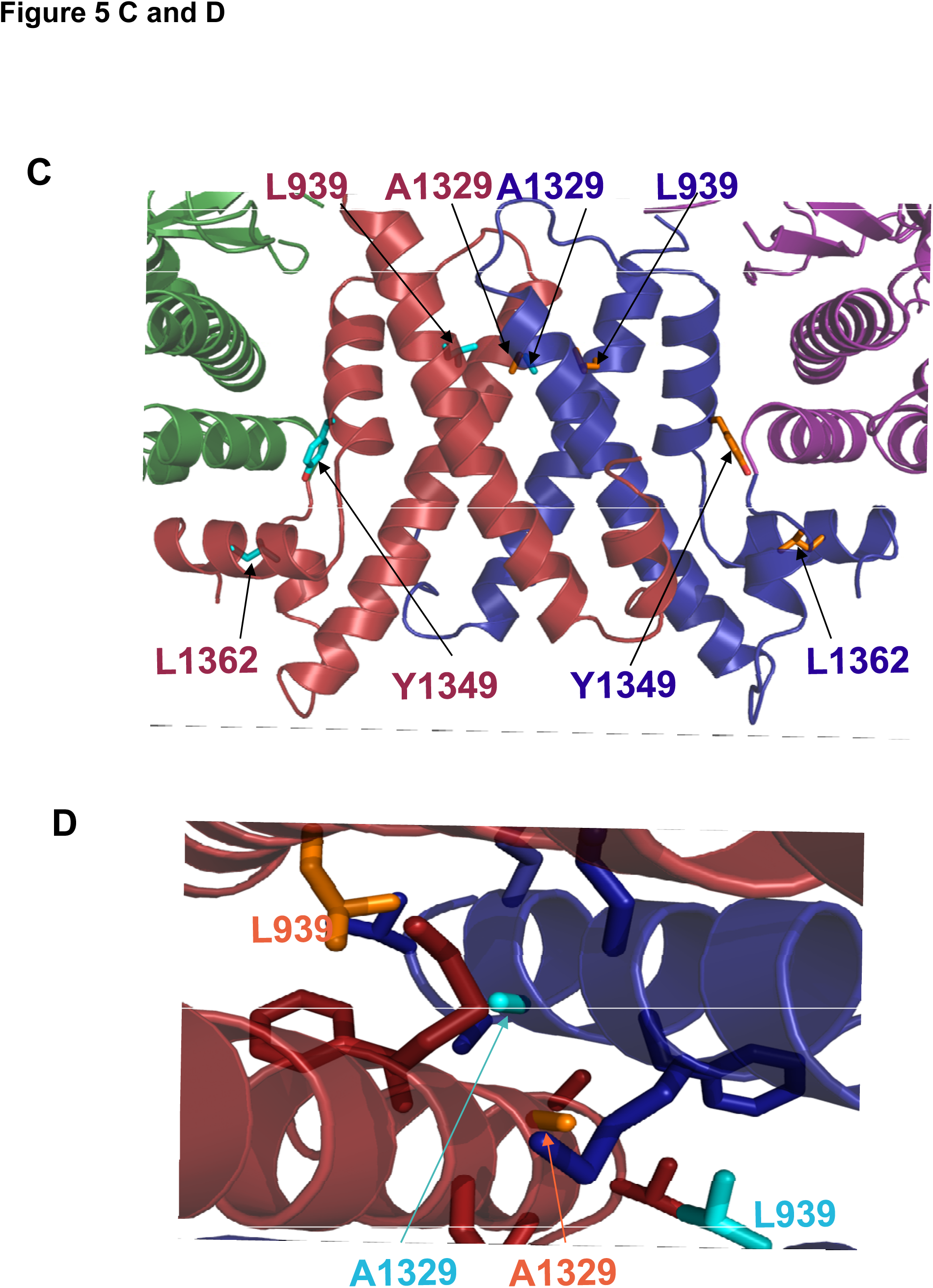
**Pragmin self-association in human cells. A.** Schematic representation of Pragmin mutants used in this study. **B.** Structure-association analysis of exogenous Pragmin in HEK293T cells. Co-immunoprecipitation of exogenous Myc-tagged wild-type or Pragmin mutants with exogenous FLAG-Pragmin (wild-type) in HEK293T cells. Is shown the levels of immunoprecipitated (IP) Myc-Pragmin and associated FLAG-Pragmin. FLAG-Pragmin levels from whole cell lysates (WCL) is shown. **C.** Zoom into the Pragmin dimer interface. The color code is as in panel 3b for the backbone while the side-chains of mutated residues are shown in wireframe in cyan and orange color. **D.** Zoom in the buried environment of alanine A1329. Neighboring side-chains are shown in wireframe and in color corresponding to their respective backbone but for A1239 and L939 which have been mutated to validate in solution the dimerization observed in the crystal structure. Picture drawn using Pymol.

### Pragmin dimerization regulates protein tyrosine phosphorylation

We next addressed the role of Pragmin dimerization on its ability to induce cellular protein tyrosine phosphorylation. Consistent with previous observations made in human CRC cells (Leroy et al, 2009), Pragmin expression in non-transformed HEK293T cells induces a substantial level of protein tyrosine phosphorylation (Fig 6A). Structure-activity analysis confirmed that the catalytic lysine K997 was largely dispensable for this molecular response. In contrast, it primarily requires its main tyrosine phosphorylation site, Y391(Leroy et al, 2009)(www.phosphosite.org). Deletion of Pragmin C-terminus including the kinase domain and dimerization extensions also strongly reduced this molecular response (Fig 6A). To ascertain the role of protein dimerization, we took advantage of Pragmin A1329E that fails to self-associate and we found that this mutant displays a reduced ability to induce protein tyrosine phosphorylation (Fig 6B). Point mutations in the dimerization domains that do not impact on Pragmin dimerization, do not affect this phosphorylation process, showing specificity (Supplementary Fig S3A). We thus concluded that protein tyrosine phosphorylation induced by Pragmin is regulated by protein dimerization and tyrosine phosphorylation.

**Figure 6:**
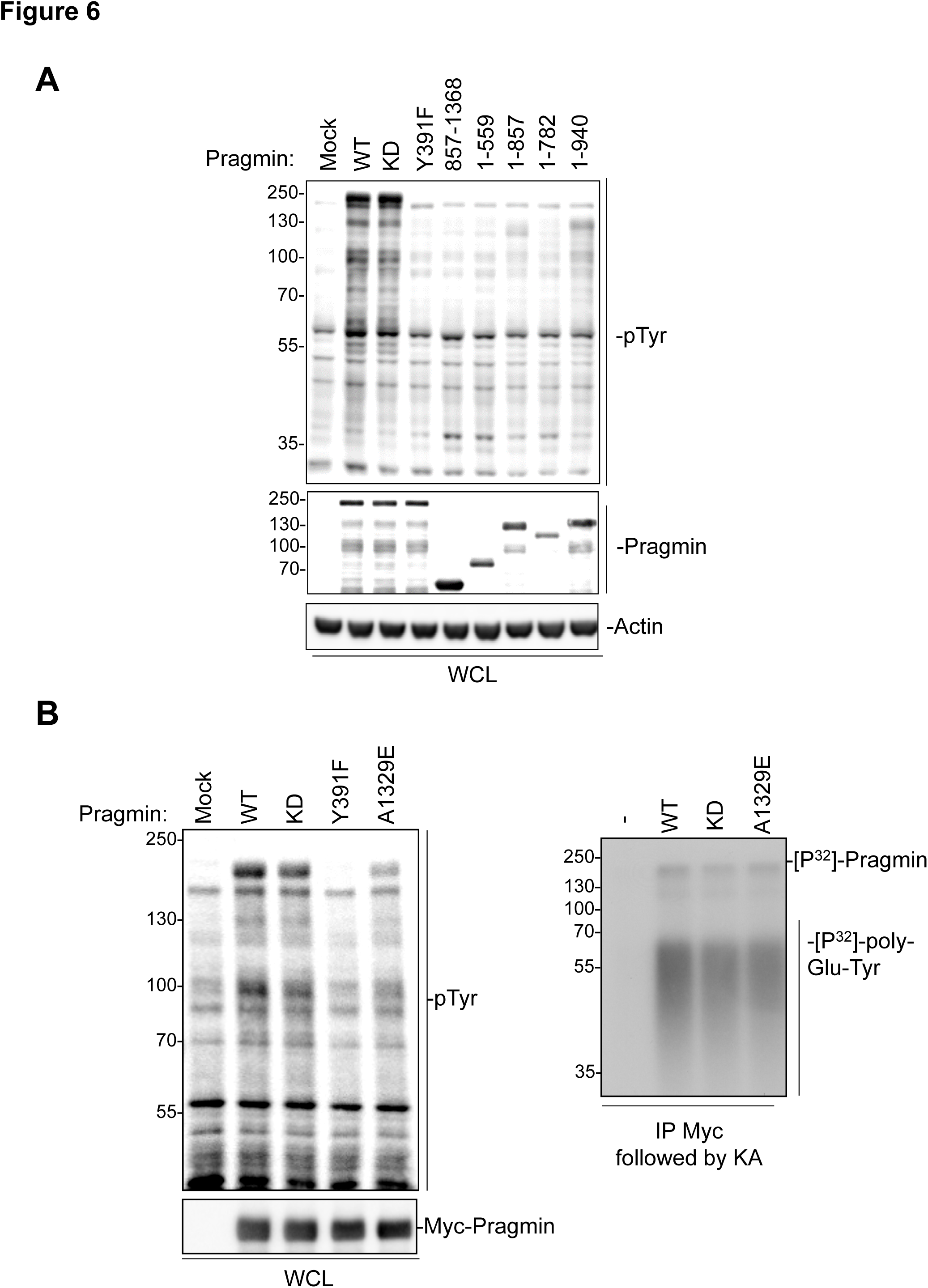
**Pragmin dimerization regulates protein tyrosine phosphorylation. A.** Exogenous Pragmin requires Its C-terminal domain to induce protein tyrosine phosphorylation in HEK293T cells. Structure-activity analysis of exogenous Pragmin in HEK293T cells. **B.** A1329E mutation reduces the protein tyrosine phosphorylation level induced by exogenous Pragmin (left) and its associated TK activity (right) in HEK293T cells. Is shown the cellular protein tyrosine phosphorylation levels from HEK293T cells transfected with indicated constructs and the in vitro TK activity present in indicated Pragmin immunoprecipitates. TK activity was measured by the level of phosphorylated poly-Glu-Tyr used as an in vitro substrate in the presence of [^32^P]-ATP ([^32^P]-poly-Glu-Tyr). The level of phosphorylated Pragmin ([^32^P]-Pragmin) is also shown.

### Self-associated Pragmin activates CSK to induce protein tyrosine phosphorylation

Since we demonstrated that Pragmin is indeed a pseudo-kinase, we hypothesized that it interacts with an active TK to induce the observed protein tyrosine phosphorylation in human cells. Consistent with this idea, we found an *in vitro* TK activity in a FLAG-Pragmin immunoprecipitate that was expressed in HEK293T cells (Fig 6C). We also noticed *in vitro* Pragmin phosphorylation in this kinase assay, indicating that this pseudo-kinase is also a direct substrate of the associated TK (Fig 6C). As expected, this activity does not require the catalytic Pragmin K997 residue. However, the A1329E Pragmin mutant associates with less TK activity, suggesting that Pragmin dimerization is required for efficient TK association and/or catalytic activation (Fig 6C and Supplementary Fig S3). No such reduced TK activity was detected on Pragmin constructs with point mutations in the dimerization domains that do not impact on Pragmin dimerization, showing specificity (Supplementary Fig S3). To identify such a TK, we next undertook a SILAC-based quantitative proteomic analysis, which allowed the identification in a semi-quantitative manner of Pragmin interactors in FLAG-Pragmin transfected HEK293T cells (Fig 7A). We reproducibly obtained more than 82 specific interactors with a mean ratio >4 and found 2 protein kinases that were prominently associated with Pragmin (mean ratio >12) including the Ser/Thr kinases AMPK and the TK CSK (Supplementary Table S2). Previous Pragmin interactors such as Src (Leroy et al, 2009) and SgK269/PEAK1 (Liu et al, 2016) were however not recovered in our SILAC analysis. The prominent interaction of Pragmin with endogenous AMPK and CSK was next confirmed by co-immunoprecipitation experiments, thus validating our SILAC analysis (Supplementary Fig S4). We also verified that, consistent with previous studies, Pragmin-CSK association primarily involves a Pragmin pY391-CSKSH2 dependent mechanism (Fig 7B) as no interaction was observed with Pragmin Y391F or any mutant that do not contain Y391(Safari et al, 2011) (Supplementary Fig S4). However, A1329E mutation did not affect Pragmin-CSK complex formation (Fig 7D), suggesting that protein dimerization is not necessary for CSK binding.

**Figure 7:**
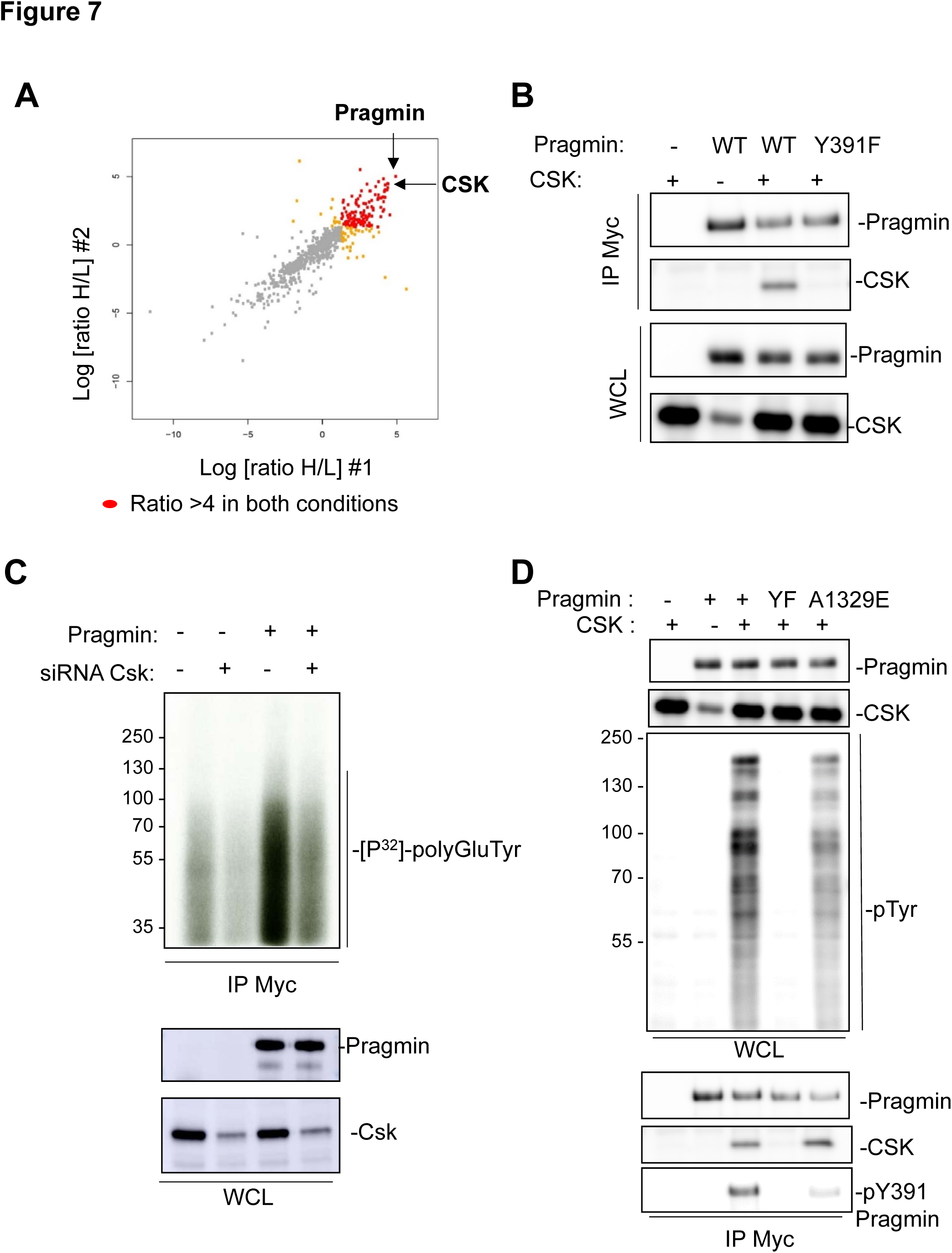
**Pragmin activates CSK to induce protein tyrosine phosphorylation. A.** SILAC analysis of exogenous Pragmin interactome in HEK293T cells reveals CSK as the main associated TK. Is shown the comparative analysis of the log of the H/L ratio of binders from two biological replicates. CSK and Pragmin are indicated. Protein with a ratio >4 in two replicates are highlighted in red and proteins with a >3 are in yellow. **B.** Biochemical analysis of exogenous Pragmin-CSK complex formation in HEK293T cells. **C.** CSK silencing drastically reduces the level of TK activity associated with exogenous Pragmin in HEK293T cells. Is shown the in vitro kinase activity ([^32^P]-poly-Glu-Tyr level) present in Pragmin immunoprecipitates from cells co-transfected with Pragmin and indicated siRNA. **D.** Pragmin dimerization stimulates CSK activity without affecting its association in HEK293T cells. Is shown the global protein tyrosine phosphorylation levels and pTyr391-Pragmin level of HEK293T cells transfected with CSK and Pragmin constructs, alone or together. The level of CSK associated with Pragmin wild-type and mutants as well as the indicated protein levels is also shown.

We next addressed whether CSK is the sought TK associated with Pragmin. We found that siRNA-mediated CSK depletion in cells expressing exogenous Pragmin resulted in a strong reduction of TK activity associated with immunopurified Pragmin (Fig 7C). Conversely, Pragmin co-expression with CSK induced a remarkable increase in protein tyrosine phosphorylation, unlike CSK or Pragmin alone (Fig 7D). This molecular response was accompanied by a strong phosphorylation of Pragmin on Tyr391 (Fig 7D), consistent with the notion that Pragmin is a novel CSK substrate (Senda et al, 2016). This increase in protein tyrosine phosphorylation was abrogated by Pragmin Y391F mutation, indicating that CSK activation primarily involves Pragmin interaction (Fig 7D). Since the CSK-Pragmin complex was reported to deregulate Src in tumor cells (Safari et al, 2011), we also examined the involvement of Src activity on this molecular process. However, cell treatment with the Src-like inhibitor SU6656 (2 µM) had a little effect on the level of protein tyrosine phosphorylation induced by CSK/Pragmin co-expression, while it strongly inhibited the one induced by Src expression, ruling out a major role for Src on this phosphorylation process (Supplementary Fig S5). We also addressed the role of Pragmin dimerization on CSK activation. We found that both the global protein tyrosine phosphorylation and the Pragmin pY391 levels induced by Pragmin/CSK co-expression was strongly diminished by Pragmin A1329E mutation (Fig 7D). Since this mutation does not affect CSK binding (Fig 7D) we concluded that Pragmin dimerization contributes to CSK catalytic activation. Finally, we addressed whether a similar mechanism operates for the main Ser/Thr protein kinase associated with Pragmin, AMPK. Like Csk, Pragmin interaction with AMPK also induced a prominent kinase activation, as determined on the pThr183 level and like CSK, AMPK activation absolutely requires Pragmin association. However, protein dimerization had not significant impact on the capacity of Pragmin to associate with, and increase AMPK activity, showing the specific role of Pragmin dimerization on CSK activation.

## DISCUSSION

Here we present a structural analysis of Pragmin signaling in human cells. First, our crystal structure of Pragmin C-terminus demonstrates that it is indeed a pseudo-kinase. Consistent with this observation, the protein kinase fold is distantly related to other kinases (RMSD ~ 3 A) in agreement with an important sequence divergence. In addition, a high sequence variability of the Hanks motifs is observed among Pragmin/Sgk269/PEAK1 sequences and the newly added members (e.g.: human C19Orf35). Finally, a unique «inhibitory triad» tightly surrounds the putative catalytic cleft suggesting inactivity in agreement with the tightly closed conformation of the crystal structure. The absence of kinase activity is further supported by the absence of any detectable adenosine or ATP binding in Pragmin as monitored by various techniques (thermal shift assay, covalent labeling,…) and the absence of significant biological effect of the K997A mutation.

Our SILAC-based interactomic analysis next identified CSK as the main TK associated with Pragmin and clarifies the mechanism by which Pragmin induces protein tyrosine phosphorylation. CSK is a TK known to essentially negatively regulate Src family tyrosine kinases (SFK) by direct phosphorylation on their conserved negative regulatory tyrosine (Sirvent et al, 2012a). Therefore, by interacting with CSK in the cytoplasm, Pragmin may prevent CSK membrane translocation for efficient membrane-associated SFK inhibition, leading to an increased SFK-dependent protein phosphorylation, as suggested in some tumor cells (Safari et al, 2011). However, our results indicate that protein tyrosine phosphorylation induced by the Pragmin-CSK complex is largely independent from SFK. Rather Pragmin association may dictate the capacity of CSK to phosphorylate novel substrates, including Pragmin itself. This notion is largely consistent with previous reports showing that CSK can signal independently from Src. For instance, CSK plays a critical role in mediating G protein signals to actin cytoskeletal reorganization (Lowry et al, 2002); eukaryotic elongation factor 2 (eEF2) phosphorylation induced by CSK promotes proteolytic cleavage and nuclear translocation of eEF2 to induce cell transformation (Yao et al, 2014); and tyrosine phosphorylation of CD45 phosphotyrosine phosphatase by CSK modulates immune signaling (Autero et al, 1994). Besides, regulation of Src-like activity during evolution, such as in the unicellular choanoflagellate *Monosiga ovata* was not coupled to CSK, suggesting an original Src-independent function for CSK (Segawa et al, 2006).

Our structural analysis also reveals a unique dimerization motif with no structural similarity with other known helical bundles. This motif regulates Pragmin self-association in human cells, which is consistent with a previous study showing a role for the pseudo-kinase N-terminal extension in protein dimerization (Liu et al, 2016). Interestingly, the dimerization interface is huge, suggesting high stability in solution. Indeed, simple mutations failed to disrupt the dimerization except the most drastic one (A1329E), probably due to a combination of several molecular effects (cumulative charge repulsion and van der Waals clashes). Our biochemical analysis reveals that this dimerization module defines an original and additional important mechanism of Pragmin signaling to induce CSK activation. CSK/Pragmin complex formation primarily involves CSK-SH2 domain and a (EPIYA) tyrosine-phosphorylation motif present in Pragmin sequence, which is used by bacterial proteins to subvert host cell signaling and maximize bacterial infection (Safari et al, 2011). However, such pTyr-SH2 dependent interaction leads to a moderate increase in CSK catalytic activity *in vitro* (Takeuchi et al, 2000), which does not match the robust capacity of Pragmin to dramatically increase CSK activity, as observed both *in vitro* (Senda et al, 2016) and in human cells. Our results indicates that Pragmin dimerization may lead to maximal CSK activation, as a Pragmin monomeric mutant displays a much reduced capacity to activate CSK while not affecting CSK binding. The molecular mechanism underlying this catalytic process is however unknown. Since side-by-side dimerization defines a general mechanism for kinase activation, we would propose a model in which the pseudo-kinase domain of Pragmin stabilizes the protein kinase domain of CSK in a fully active conformation by direct interaction. However, we did not detect any interaction between these domains in co-immunoprecipitation experiments. While our data does not favor this model, we cannot rule out any low affinity interaction, which was not detected by standard biochemical methods. Another model would imply that that self-associated Pragmin interacts with two molecules of CSK in the same complex and favors CSK homo-dimerization in a fully opened and active conformation. Clearly, further experiments will be needed to validate any of these models.

According to our structural characterization, Pragmin and the related proteins (SgK269/PEAK1 and C19orf35) constitute a new family of scaffolding pseudo-kinases. We believe that the important molecular insights obtained from our structural analysis may apply to the other members of the family. Firstly, our results predict that SgK269/PEAK1 is also a pseudo-kinase, therefore the reported capacity of SgK269/PEAK1 to activate protein tyrosine phosphorylation (Wang et al, 2010) may involve the association of an active TK, possibly a member of the Src Family Kinases (SFK) (Croucher et al, 2013; Kelber et al, 2012). It will be interesting in this context to see whether C19Orf35 also induces protein phosphorylation and if yes, what the nature of the associated active protein kinase is. Secondly, their N- and C-terminal extensions may form a dimerization module with similar regulatory functions. This prediction is consistent with a previous study showing a role for the pseudo-kinase N-terminal extension in SgK269/PEAK1 in protein dimerization (Liu et al, 2016). Like Pragmin, protein dimerization may regulate the activity of the associated protein kinase. Finally, the dimerization modules may also induce heterotypic association as experimentally observed for Pragmin and SgK269/PEAK1 (Liu et al, 2016). The level of sequence conservation at the interface within the family would also predict heterotypic association with C19Orf35. The resulting complexes may modulate the strength and the nature of signaling induced by these pseudo-kinases, as reported for Pragmin/SgK269 heterotypic association. In this regard, it will be important to test whether heterotypic Pragmin association reduces CSK signaling and promote Notch-dependent transcription.

Finally, our structural analysis may have a significant impact in our understanding of the oncogenic role of pseudo-kinases, such Pragmin in human cancer. Our observations support the idea that Pragmin induces an unexpected tumoral activity of CSK in human cancer, such as CRC. While CSK displays anti-oncogenic activity in some cancers by inactivation of SFK (Masaki et al, 1999; Okada, 2012; Sirvent et al, 2010), this mechanism is impaired in CRC due to downregulation of the CSK transmembrane binder CBP/PAG (Sirvent et al, 2010). Rather, the protein CSK is strongly overexpressed in CRC along with aberrant expression of SFK activity and patients display CSK autoantibodies, which defines a novel biomarker of the disease (Oneyama et al, 2008; Sirvent et al, 2010). CSK overexpression also increases growth of advanced CRC cells (Sirvent et al, 2010) and induces migration of additional epithelial tumor cells in a Pragmin-dependent manner (Senda et al, 2016). Conversely, Pragmin invasive activity requires its CSK-binding site(Leroy et al, 2009). Therefore Pragmin upregulation combined with deregulation of upstream TKs occurring during tumor progression may turn CSK oncogenic in CRC. Targeting this Pragmin signaling may therefore be of therapeutic interest in these advanced tumors.

## METHODS

### Sequence-structure analysis

Fold-recognition were performed using the server @TOME-2 (Pons & Labesse, 2009). Domain boundaries of the predicted protein kinase domain were refined manually using ViTO, an editor of sequence-structure alignment (Catherinot & Labesse, 2004). Secondary structure predictions were performed using JPRED3 (Cole et al, 2008). Subsequently, partial models built using @TOME-2 from the best matching templates were gathered and superposed using the software MATT (Menke et al, 2008) and used an ensemble for molecular replacement (see below).

### Antibodies and reagents

Anti-Myc 9B11 (#2276; 1:1,000), anti-Biotin D5A7 (#5597; 1:1,000), anti-HES-1 D6P2U (#11988, 1:1,000), anti-AMPK D63G4 antibodies (#5832; 1:1,000), anti-rabbit IgG-HRP (#7074S, 1:5000) and anti-mouse IgG-HRP (#7076S, 1:5,000) were from Cell Signaling; anti-Actin AC-15 (#A5441, 1:5,000) and anti-FLAG antibody (#F7425, 1:1,000) and anti-FLAG M2 Affinity Gel (#A2220) from Sigma-Aldrich; anti-Myc-Agarose Beads (9E10) (#631208) was from Clontech; anti-mouse Alexa-488 (#A11001; 1:1,000), anti-rabbit Alexa-594 (#A11012; 1:1,000) and Rhodamine Phalloidin (#R415, 1:1000) from Life technologies; anti-Tubulin (a gift from N. Morin; CRBM) (1:2,000), anti-pTyr 4G10 (a gift from P. Mangeat, CRBM)(1:50); anti-CSK (1:1,000) and anti-SFK CST-1 antibodies (1:1,000) were described in(Benistant et al, 2001; Sirvent et al, 2010). Anti-pY391 Pragmin rabbit antibody was raised against AVQPEPIpYAESAKRK peptide and affinity-purified by Eurogentec.

### Plasmids constructions

The cDNA coding amino acids 906-1368 of Rattus norvegicus Pragmin protein (Pragmin-C) was cloned into pET28b (Novagen) vector between NdeI and XhoI sites to give the pET 28b-Pragmin906WT vector. In this construct, the protein was fused with an N-terminal thrombin-cleavable His6 tag. Pragmin D978N, Y981F and Q1021E mutations were generated by PCR-based site-directed mutagenesis (QuikChange, Agilent) with the pET28b-Pragmin906WT vector as template. pCMV-Myc-Pragmin (a gift of Dr Negishi, Kioto University), pSGT-Src (Chicken) WT and KD (K295M), pcDNA3-CSK (Human) and pMX-CESAR Myc-Pragmin were respectively described in(Leroy et al, 2009; Roche et al, 1995; Sirvent et al, 2010; Tanaka et al, 2006). The FLAG-Pragmin construct was generated by digesting pCMV-Myc-Pragmin with EcoRI and NotI and subcloned into the pCMV5-Flag vector. Control siRNA or CSK siRNA (ON-TARGET plus siRNA #03110-10) were purchased from Dharmacon. Pragmin mutant constructs were generated from pCMV-Myc-Pragmin by PCR using the QuickChange II Site-Directed Mutagenesis Kit (Agilent) and the corresponding oligonucleotides:

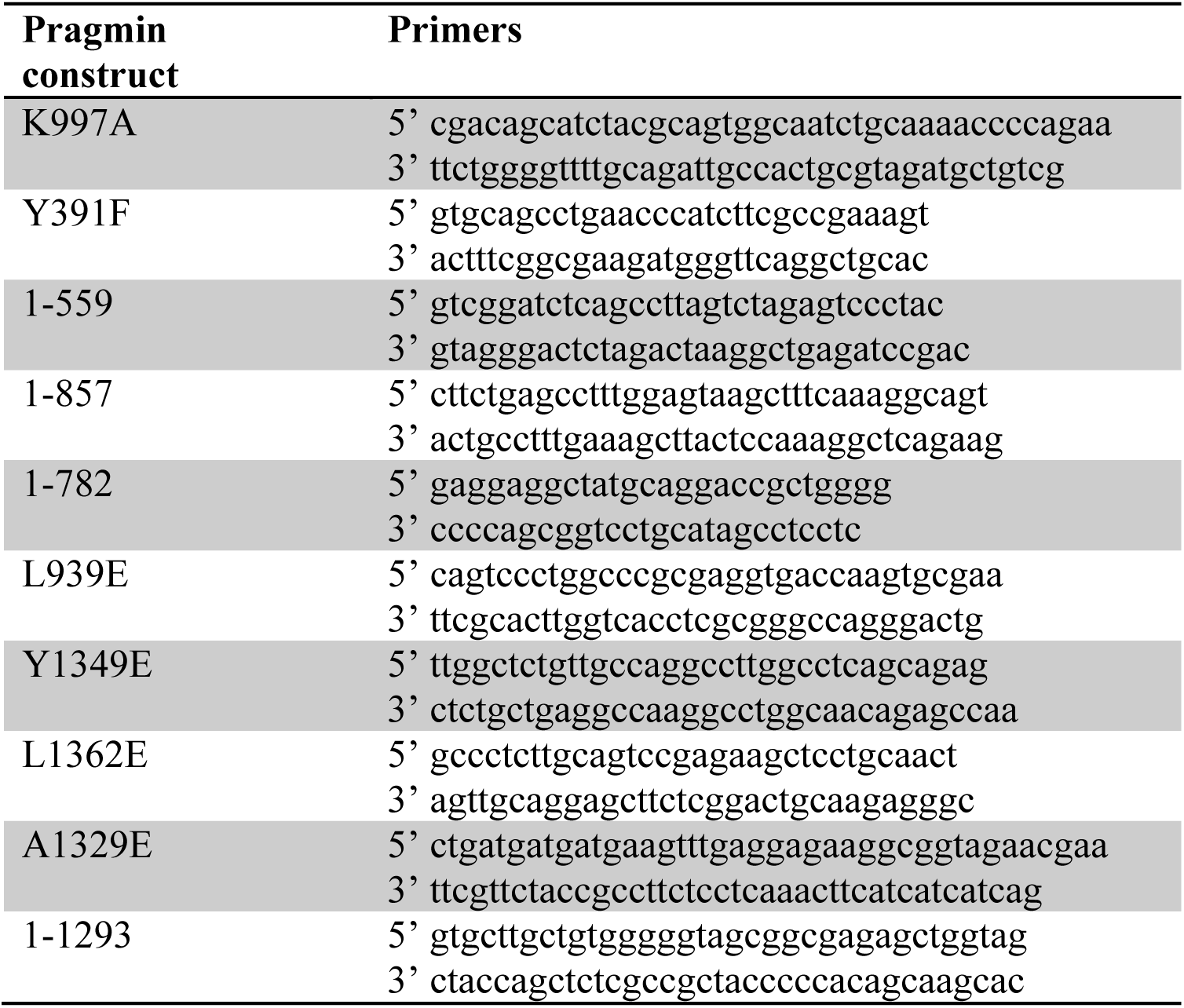

### Proteins expressions and purifications

Wild-type and mutants constructs were transformed into Escherichia coli RosettaTM(DE3) strain. Transformed cells were grown at 37°C in LB medium supplemented with 50 µg/ml kanamycin. When turbidity reached A600 = 0,6, the temperature was switch to 16°C, Pragmin expression was induced overnight by addition of 0.5 mM IPTG. Cells were harvested by 20 min centrifugation at 6000g at 4°C. The pellet was resuspended in Tris-HCl 20 mM pH 7.5, sodium chloride 0.3 M and 2 mM 2-mercaptoethanol (buffer A) and stored at −80°C. Cells were supplemented with a Complete® EDTA free tablet (Roche), lysed by sonication, and insoluble proteins and cell debris were sedimented by centrifugation at 40,000g at 4°C for 30 min. Supernatant was supplemented with imidazole to 10 mM final concentration, filtered through 0.45-μm filters and loaded onto affinity column (5 ml His Trap FF, GE Lifescience) that was equilibrated with buffer B (buffer A containing 10 mM imidazole). Column was washed with 20 column volume of buffer B and proteins were eluted with a linear 0–100% gradient of buffer C (buffer A containing 0.5-M imidazole). The peak fractions were analyzed by SDS-PAGE. Fractions containing Pragmin-C were pooled and dialyzed overnight at 4°C against a conservation buffer containing sodium citrate 50mM pH 5,5; sodium chloride 0.2 M; DTT 5mM (buffer D). Pragmin-C was concentrated to 1.2-1.5 mg/ml and the 6His-tag was eliminated by Thrombin cleavage (ratio protease/protein 2U/mg) overnight at 4°C. Cleaved proteins were injected onto a gel filtration column (Superdex 75 Hiload 16/60 GE Lifescience) equilibrated with buffer D. The peak fractions were analyzed by SDS-PAGE and the fractions containing pure Pragmin-C were pooled and concentrated to 7 mg/ml with Vivaspin centrifuge concentrator (Sartorius stedim biotech) after which, crystallization screening was immediately carried out.

### Crystallization

Initial crystallization conditions of Pragmin-C were found by using a sitting-drop based and sparse-matrix screening strategy. Four hits were detected, one corresponding to a salt crystal (mother liquor containing 1.3 M Na/K tartrate, pH 5.6) and the three other to a protein crystal (all containing ammonium sulphate from 0.2 M to 1 M), as monitored by in situ X-ray diffraction on BM14. The best diffraction pattern (up to a 4.5 Å resolution) was obtained for crystal grown in ammonium sulphate 1 M, potassium chloride 1 M and 0.1 M HEPES pH 7.0). After stepwise refinement, well-diffracting crystals were obtained by mixing 1 µl of the protein solution (concentration 7.5 mg/ml) with an equal volume of crystallization buffer (ammonium sulphate 1M, lithium citrate 0.2 M pH 5.0 and 12.5 mM EDTA), equilibrated over 0.4 ml of the same buffer.

### Crystallographic Studies

X-ray diffraction data sets were collected from frozen single crystals at the European Synchrotron Radiation Facility (Grenoble, France, beamlines BM14, MASSIF-1, ID23-2 and ID23-1). Data were processed and scaled with the program suites installed at the ESRF (Monaco et al, 2013). Anisotropy was corrected after scaling (Strong et al, 2006). The structure was solved by molecular replacement using AutoMR procedure of the software PHENIX (Adams et al, 2010) using a dataset at a 3.0-Å resolution and an ensemble-based model built using @TOME-2 (see above). Improvement of the solution was obtained by automatic model building with the program AutoBuild (Adams et al, 2010). Subsequently, Iterative model rebuilding and refinement was performed first by using the program COOT (Emsley et al, 2010) and the program PHENIX (Adams et al, 2010), using a translation/libration/screw model against the native data set at 2.77-Å resolution (Supplementary Table S1). In the final model, various segments of the protein were not clearly visible in the electron density map and mainly correspond to loops variable in sequence and length among Pragmin orthologues. Figures of ligands and corresponding electronic density were generated using PyMOL (http://pymol.sourceforge.net).

### Biophysical characterization

Integrity of the wild-type and mutated recombinant proteins were checked using far-UV circular dichroism (data not shown). Thermal Shift Assay (hereafter TSA) assays were set up using an Mx3005P quantitative PCR instrument (Stratagene) as previously described(Niesen et al, 2007) using SYPRO orange (Sigma, n° S5692) as a dye. SYAL filter corresponding to the optimal excitation (492 nm) and emission (610 nm) wavelengths for SYPRO orange dye was used with the experimental mode "SYBR Green (With Dissociation Curve)". The temperature was increased by 1°C each cycle over a temperature range of 25-95 °C. Assay reactions were performed in 96-well white PCR plates (ThermoScientific, Thermo-Fast low Profile, N° 39365), and wells were capped using optical tape (Bio-Rad, n° 223-9444). Data were exported and Tm determined thanks to fitting to the Boltzmann equation (GraphPad Prism5). First, to search for optimal stabilization conditions and make protein sample freezable, an in-house buffer (pH 3,1 to 9,5) and additives screen was used. Pure protein purified in buffer E (HEPES pH 7.5 50mM, potassium citrate 0.15 M, DTT 5mM) was mixed 1:1 with SYPRO orange dye. 10 µL was dispensed in the wells of the PCR plate, and mixed with equal volume of each screening solution (1.6 µM final protein concentration). The reference Tm (with no additive) was 57.3°C. The best stabilizing buffer was sodium citrate pH 5.3 (delta Tm = +1.6°C), and was further improved with addition of 0.5 M sodium chloride (delta Tm = +3.3°C). Wild-type enzyme and its variants D978N, Q1021E and Y981F were also assayed with the buffer screen in the same conditions as above except for final protein concentration (2 µM) and starting buffer (lithium citrate 0.1 M pH 5.2, DTT 5 mM). The reference Tm for each variants were 55.7°C, 53.4°C, 55.5°C and 49.8°C, respectively. The stability of the three mutants shows same global trends as the wild-type enzyme, including a generally favorable effect of ammonium sulphate (0.2 M, up to 2.4°C delta Tm). The impact of ADP, ATP and MgATP was monitored at two final concentrations (5 and 10 mM). The ligands dissolved in water at 100 mM were diluted in protein buffer and 10 µL of the resulting solutions were added to the 10 µL of protein-dye mixture as previously. None showed any significant stabilizing effect on any of the four variants tested.

### Cell culture, transfections and retroviral infections

HEK293T (ATCC, Rockville, MD, USA) were cultured in Dulbecco’s Modified Eagle’s Medium (DMEM) Glutamax supplemented with 10% fetal calf serum (FCS), 100 U/ml penicillin and 100 mg/ml streptomycin at 37°C and 5% CO2 in a humidified incubator. Transfections in HEK293T were performed using jetPEI (Polyplus-transfection) and XtremeGENE HP transfection reagent (Roche) respectively, following the manufacturer's instructions. For siRNA transfection, 250,000 cells were seeded in six-well plate and transfected 24h later using 30 pmol siRNA and 9 µL Lipofectamine® RNAiMAX (Life Technology) for 48h. For inhibitor assays, cells were treated with 2 µM SU6656 (Merck-Millipore) for 2-3h.

### Biochemistry

Immunoprecipitations (IP) and Western blotting (WB) were performed as described in(Sirvent et al, 2010). Briefly, cells were rinsed twice in PBS and scraped in 2x lysis buffer containing 1% Triton X-100, 10 mM Tris-HCl (pH 7.5), 150 mM NaCl, 5 mM EDTA, 75U/ml aprotinin, and 1 mM vanadate. Proteins were separated on SDS-PAGE gels and transferred onto Immobilon membranes (Millipore Molsheim, France). Detection was performed using the ECL System (GE Healthcare). Optimal exposure times of membranes were used. In vitro kinase assay was performed as described in (Sirvent et al, 2010) using 10 µg poly-Glu-Tyr (Sigma) as a substrate in the presence of 0.1 mCi [^32^P]-ATP (Amersham) and incubated at 30°C for 10 min. ActivX Desthiobiotin-ATP binding was performed following the manufacturer’s instructions (Pierce). Briefly, immunoprecipited proteins were incubated with MgCl2 (20 µM, 1 min at RT), then with Desthiobiotin-ATP (20 µM, 30 min, 30°C). After washing, proteins were loaded on SDS-PAGE gel and Desthiobiotin labeled proteins were detected by anti-biotin antibody.

**SILAC analysis.** SILAC analysis was performed essentially as previously described in(Naudin et al, 2014). HEK293T cells were cultured in SILAC DMEM (Pierce) without Lysine (Lys) and Arginine (Arg) and supplemented with 4 mM L-glutamine, 10% dialyzed FBS (Invitrogen), 0.084 g l^-1^ Arg and 0.146 g l^-1^ Lys. Heavy (^13^C615N4-Arg and ^13^C615N2-Lys, from EurisoTop) or unlabeled amino acids (light Arg and Lys, from Sigma Aldrich) were used. After 3 weeks of metabolic labelling, cells were transfected with FLAG-Pragmin (heavy conditions) or empty vector (light conditions) for 48 hrs and lysed in lysis buffer. Cell-lysates (30 mg protein) were combined and incubated overnight with anti-FLAG antibody coupled to protein G bound agarose beads to avoid the presence of immunoglobulin heavy chains in samples. Purified proteins were separated on SDS-PAGE gels and trypsin-digested samples (1 µl) obtained from cut gel slices were analyzed using a Qexactive system coupled with a RSLC-U3000 nano HPLC apparatus. Desalting and pre-concentration of the samples were performed on-line on a Pepmap® precolumn (0.3 mm x 10 mm). A gradient consisting of 0-30% B for 65 min, 30-50% in 15 min and 90% B for 10 min (A = 0.1% formic acid, 2% acetonitrile in water; B = 0.1 % formic acid in 80% acetonitrile) at 300 nl/min was used to elute peptides from the capillary (0.075 mm x 150 mm) reverse-phase column (Pepmap®, Dionex), fitted with an uncoated silica PicoTip Emitter (NewOjective, Woburn, USA). Spectra were acquired with the instrument operating in the information-dependent acquisition mode throughout the HPLC gradient. Survey scans were acquired in the Orbitrap system with resolution set at a value of 70,000. Up to ten of the most intense ions per cycle were fragmented and analyzed using a resolution of 17,500. Peptide fragmentation was performed using nitrogen gas on the most abundant and at least doubly charged ions detected in the initial MS scan and an active exclusion time of 45 s. Analysis was performed using the MaxQuant software (version 1.5.0.0). All MS/MS spectra were searched using Andromeda against a decoy database consisting of a combination of Homo sapiens CPS databases (release November 2014 www.uniprot.org), Pragmin sequence and 250 classical contaminants, containing forward and reverse entities. A maximum of 2 mis-cleavages were allowed. The search was performed allowing the following variable modifications: Oxidation (M) and Phospho (STY). FDR was set at 0.01 for peptides and proteins and the minimal peptide length at 7.

## Acknowledgements

We thanks the Montpellier RIO Imaging platform and the functional Proteomic Platform of Montpellier for imaging and proteomic analyses. This work was supported by the French Infrastructure for Integrated Structural Biology (FRISBI) ANR-10-INBS-05, INCA (PLBIO-2011-150), La Ligue Contre le Cancer (équipe labellisée LIGUE2014 and 2017), FRM, Montpellier SIRIC Grant «INCa-DGOS-Inserm 6045», CNRS and the University of Montpellier. SR is an INSERM investigator.

## Accession codes

Coordinates and structure factors for rat PRAGMIN-906-1368 have been deposited in the Protein Data Bank under accession code PDB.

## Contributions

GL and SR designed the experiments and wrote the manuscript. VS, FA and AF conducted protein production and directed mutagenesis. MG performed thermostability and covalent attachment assays. MG and GL performed crystallization, X-ray data collection and structure refinement. Sequence-structure alignment and structural analysis was performed by JLP and GL. CL, CB, VS IM and CK performed biochemical and cell biological analyses. CL and SU performed SILAC analyses. Correspondence to: GL and SR.

## Supplementary Figures

**Supplementary Figure S1: Sequence alignment of Pramin orthologues and paralogues.** Sequences of Pragmin/SGK223 and PEAK1/SGK269 proteins from rat, human and xenopus were extracted for UNIPROT (http://www.uniprot.org/uniprot/) and named PRAG1 or PEAK1 followed by the species name following UNIPROT nomenclature. CS035 from human was also extracted from UNIPROT while the two other orthologues (CS035 from *Loxodonta Africana* and *Monodelphis domestica* are named herein cs035_loxaf and cs035_mondo, respectively) were from Genpept. Sequence alignment was edited using ViTO to optimize positions of insertions/deletions. Secondary structure were computed from the crystal structure of Pragmin and numbered following the the protein kinase nomenclature for the pseudo-kinase domain. Hank's motifs are labelled using roman numbering under the alignment. Domain delimitation are shown by right and left arrows in black color. Residues not visible in the electron density from either monomer are underlined using grey and filled circles. Mutated residues involved in the “inhibitory triad” and the dimerization interface are highlighted by black stars and filled circle, respectively. The figure was produced using ESPRIPT (Robert & Gouet, 2014).

**Supplementary Figure S2: UV-CD spectra of wild-type protein and mutants in the putative ATP-binding site.** UV-CD were recorded a 0.7 mg/ml in 100 mM lithium citrate buffer at pH 5.2 with a path length 0.2 mm on a AppliedPhotoPhysics Chirascan.

**Supplementary Figure S3: Effect of point mutations in the dimerization domains on Pragmin-induced protein tyrosine phosphorylation. A.** Protein tyrosine phosphorylation level induced by exogenous Pragmin (wild-type and indicated mutants) and the level of its associated TK activity (**B)** in HEK293T cells. In vitro TK activity present in indicated immunoprecipitates was measured by the level of phosphorylated poly-Glu-Tyr used as an in vitro substrate in the presence of [^32^P]-ATP ([^32^P]-poly-Glu-Tyr). The level of associated TK activity (relative to the activity obtained in Pragmin IP) is shown; (mean ± SEM; n=4).

**Supplementary Figure S4: mutagenesis analysis of Pragmin association with CSK and AMPK A.** Schematic representation of Pragmin mutants used in this study. co-immunoprecipitaiton of exogenous Pragmin mutants with endogenous CSK **(B)** and AMPK **(C)** in HEK293T cells. The level of AMPK activity (pThr183) is also shown. *represents the IgG Hc.

**Supplementary Figure S5: Src is not involved in Pragmin-induced CSK activation** Protein tyrosine phosphorylation induced by Pragmin co-expression with CSK in HEK293T cells (as shown) is poorly affected by cell treatment with the Src inhibitor SU6656 (2 µM) while is strongly reduces protein tyrosine phosphorylation induced by Src expression.

**Supplementary Table S1:** **Data collection, phasing, and refinement statistics of Pragmin C-terminus crystallography.**

|  |  |
| --- | --- |
| **Data collection** |  |
| Space group | C 1 2 1 |
| Cell dimension<br>a, b, c (Å) | 193.43 58.92 110.64 |
| $\beta$ (°) | 118.48 |
| No. molecules in a.u | 2 |
| Wavelength (Å) | 0.97294 |
| Collection temperature (K) | 100 |
| Resolution range (Å) <sup>1</sup> | 55 - 2.77 (2.87 - 2.77) |
| Rmerge (%) <sup>1-2</sup> | 0.052(0.614) |
| CC1/2 | 0.999(0.868) |
| Mean I/σI <sup>1</sup> | 12.2(2.1) |
| Completeness (%) <sup>1</sup> | 99.2(99.9) |
| Redundancy <sup>1</sup> | 4.1(4.2) |
| B-wilson | 56.80 |
| **Refinement** |  |
| Resolution (Å) | 2.77 |
| No. Reflections | 25024 |
| Rwork/Rfree (%) <sup>3-4</sup> | 0.1905 / 0.2428 |
| No. Atoms |  |
| Protein | 5641 |
| Water | 69 |
| B-factors (Å<sup>2</sup>) |  |
| Protein | 63.3 |
| Water | 51.2 |
| R.m.s deviations <sup>5</sup> |  |
| Bond lengths (Å) | 0.009 |
| Bond angles (°) | 1.17 |
| Ramachandran favored (%) | 93.4 |
| Ramachandran allowed (%) | 5.1 |
| Ramachandran outliers (%) | 1.5 |
| Rotamer outliers (%) | 5.5 |
| all-atom clashscore | 9.79 |
| **PDB id** |  |

**Supplementary Table S2: SILAC analysis of Pragmin interactome in HEK293T cells**

